## Supplemental Methods and Text for "Early epigenetic influence on EGR1, KLF2 and KLF4 transcriptional network associates with alteration of HSCs quiescence"

#### Supplementary information

**Supplemental Figure 1: Methylation data processing.** A heatmap is shown displaying the -$\log_{10}$ *p*-values of linear regression for top ranking principal components for each known covariate. The color keys correspond to numeric values for each covariate, with red indicating greatest significance.

**Supplemental Figure 2: HSPC subpopulations analysis** (A) UMAP representing subpopulation clusters within each HSPC lineages. (B) Dot plot representing key markers used to annotate subpopulations. LT-HSC, long-term hematopoietic stem cell; HSC, hematopoietic stem cell; MPP, multipotent progenitor; LMPP, lymphoid-primed multipotent progenitor; CLP, common lymphoid progenitor; Ery, Erythroid; EMP, erythron-myeloid progenitor; ErP, erythroid progenitor; Mk/Er, megakaryocyte and erythrocyte; GMP, granulocyte-monocyte progenitor; DC, dendritic cell; cycle, in G2/M phase.

**Supplemental Figure 3: DEG analysis.** (A) Volcano plot representing DEG comparing LGA vs CTRL considering all lineage. Differentially expressed genes with adjusted *p*-value <0.05 and $|\log_2 FC|$ >0.5 are shown in red. (B) Bar plot representing the distribution across lineage for DEGs. (C) Volcano plot representing DEG analysis at cell population level comparing LGA vs CTRL. Differentially expressed genes with adjusted *p*-value <0.05 and |fold change| >0.5 are shown in red. LT-HSC, long-term hematopoietic stem cell; HSC, hematopoietic stem cell; MPP, multipotent progenitor;

LMPP, lymphoid-primed multipotent progenitors; Erythro-Mas, erythroid and mast precursor cell.

**Supplemental Figure 4: ATAC-seq data processing.** (A) UMAP representing subpopulation clusters based on chromatin accessibility. Clusters are associated to lineage based on overlap with annotations through label transfer from transcriptomic data. (B) Dot plot representing the top enriched transcription motifs within lineage specific peaks. Lineages no represented do not have lineage specific peaks. (C) Volcano plot representing differentially accessible peaks (Down/Up peaks). Differentially open peaks with adjusted *p*-value <0.001 and $|log_2FC|$ >0.25 are shown in red. (D) Dot plot representing enrichment for transcription factor motif within Up peaks identified comparing chromatin accessibility between LGA and CTRL.

**Supplemental Figure 5: TFs and pseudotime lineage specific characterization.** (A) Heatmap representing the lineage specific regulon activity highlighting lineage specific TF. LT-HSC, long-term hematopoietic stem cell; HSC, hematopoietic stem cell; MPP, multipotent progenitor; LMPP, lymphoid-primed multipotent progenitors; Erythro-Mas, erythroid and mast precursor cell. Regulon including lower confidence targets are identified with the “e” suffix. (B) Boxplots of the pseudotime distribution across lineage demonstrating a positive correlation between differentiation process and pseudotimes.

#### Supplemental Methods

**Sample collection.** Cord blood from neonates was the source of material for this study. Biological samples and clinical information were collected from consenting women who delivered healthy infants without any anomalies or dysmorphic features and following an uncomplicated intrapartum course, without evidence of fetal distress (normal Apgar scores and cord blood gases without acidemia). The groups were comprised of infants with appropriate growth (CTRL) or large for gestational age (LGA) neonates (matched for gestational age at delivery and sex). Both birth weight and ponderal index (a measurement of neonatal weight relative to length) were used to identify case and control subjects. LGA were defined by birth weight and ponderal index values greater than the $90^{th}$ percentile for gestational age and sex. Control infants had normal parameters (between $10^{th}$ and $90^{th}$ percentiles) for both birth weight and ponderal index. Maternal and infant characteristics are shown in **Supplemental Table 7**.

**Isolation of CD34+ HSPCs.** CD34+ cells, which constitute approximately 1% of nucleated blood cells in umbilical cord blood, were isolated from the cord blood specimen using an immunomagnetic separation technique. Mononuclear cells were separated using PrepaCyte-WBC following which CD34+ cells were obtained by positive immunomagnetic bead selection, using the AutoMACS Separator (Miltenyi Biotech). This resulted in the isolation of cells with 95% purity. We cryopreserved the purified cells in 10% dimethyl sulphoxide using controlled rate freezing.

**Genome-wide DNA methylation assay.** The HELP-tagging assay was performed after isolation of genomic DNA from frozen CD34+ HSPCs, digested to completion by either HpaII or MspI. The digested DNA was ligated to two custom adapters containing

Illumina adapter sequences, an EcoP15I recognition site and the T7 promoter sequence. Using EcoP15I, we isolated sequence tags flanking the sites digested by each enzyme, methylation-sensitive HpaII or methylation-insensitive MspI, followed by massively parallel sequencing of the resulting libraries (Illumina Technology). HpaII profiles were obtained for each sample, calculating methylation scores using a previously generated MspI human reference.

**Single-cell RNA sequencing libraries preparation.** Each cryopreserved CD34+ cells from each sample were thawed in a water bath at 37°C 1min before to be resuspended in 10 ml of pre-heated medium. Cell suspensions were filtered with a MACS pre-separation filter 20 µm and centrifuged 5min at 300g. Cell pellets were resuspended in Deionized Phosphate Buffer Saline 1X (DPBS, GIBCO™, Fisher Scientific 11590476) with 0.04% Bovine serum albumin (BSA) for counting on a Corning Cytosmart cell counter by Trypan blue (Trypan Blue solution, 11538886, Fisherscientific) counterstaining for viability check. Cell suspension was loaded on a Chromium 10x Genomics controller following the manufacturer protocol using the chromium single-cell v3 chemistry with single indexing. Specifically, single cells, reverse transcription (RT) reagents, Gel Beads containing barcoded oligonucleotides, and oil were combined on a microfluidic chip to form nanoliter-scale reaction vesicles. Within each reaction vesicle, a single cell was lysed, the Gel Bead was dissolved to free the identically barcoded RT oligonucleotides into solution, and reverse transcription of polyadenylated mRNA occurs. As a result, all cDNAs from a single cell will have the same barcode, allowing the sequencing reads to be mapped back to their single cells of origin. The gene expression library also carries an additional unique molecular identifier (UMI) to distinguish individual, captured mRNA molecules for quantification. The preparation of NGS libraries from these barcoded cDNAs was then carried out in bulk reaction. Gene expression libraries were sequenced using 100 pb paired-end reads on NovaSeq 6000 system following the manufacturer recommendations (Illumina) at a minimum depth of 25,000 reads. Following sequencing, BCL or FASTQ files can be analyzed using the Cell Ranger analysis pipeline. Cell Ranger performs sample demultiplexing, barcode processing, and counting of transcripts in single cells. Secondary analyses, such as dimensionality reduction, cell clustering, and differential gene expression, are performed through Seurat and detailed bellow.

**HTO protocol.** After cell counting and viability check, cell Hashtag (HTO) staining was used for cell stimulation following the cell-hashing protocol.[1] For each samples, cells were resuspended in 100 µl of staining buffer (DPBS BSA 2%, Tween-20 0,01%) with 10 µl of Fc blocking reagent HumanTruStainFc™ (422302, Biolegend) and incubated 15 min at 4°C. 1 µl of antibody was added (TotalSeq™-A anti-human Hashtag from 1 to 6, 0251, 0252, 0253, 0254, 0255, 0256, Biolegend) followed by a 30 min incubation at 4°C. Cells were washed 3 times in staining buffer with one filtration step by MACS pre-separation filter 20 µm (Miltenyi Biotec) to a final resuspension in DPBS 0.04%. Cell suspensions from each sample were pooled (*n*=3 to 6 per library) prior to loading on a Chromium 10x Genomics controller following the manufacturer protocol using the Chromium single-cell v3 chemistry with single indexing. Gene expression libraries preparation were performed as described above. HTO libraries preparation were performed using the adapted Biolegend protocol (https://www.biolegend.com/en-us/protocols/totalseq-a-antibodies-and-cell-hashing-with-10x-single-cell-3-reagentkit-v3-3-1-protocol). The resulting libraries were pooled at equimolar proportions with a 9 for 1 ratio for Gene expression library and HTO library respectively.

**Nuclei isolation for Single-cell ATAC sequencing**

After thawing CD34+ cells as describe in Single-cell RNA sequencing libraries preparation, nuclei were isolated based on 10X Genomics protocol for low cell input (<100k cells, Nuclei Isolation for Single Cell ATAC Sequencing CG000169-Rev D). Lysis time, optimized to isolate without damage nuclei, was set to 3min.

**Single-cell ATAC sequencing libraries preparation.** Chromatin accessibility was analyzed on a cell by-cell basis through the use of microfluidic partitioning to capture single cells and prepare barcoded, next-generation sequencing (NGS) libraries. Transposition is performed in bulk upon application of the enzyme transposase, which enters the nuclei and preferentially fragments the DNA in open regions of chromatin while adapter sequences are simultaneously added to the ends of the DNA fragments. Transposed nuclei are loaded onto a microfluidic chip, which is run in the Chromium Controller instrument. In the instrument, nuclei are partitioned individually with a single Gel Bead forming droplets, or Gel Beads-in-emulsion (GEMs). Each Gel Bead contains oligonucleotides with a unique 16 base pair 10x Barcode sequence and matching adapter sequence that enables attachment of transposed DNA fragments for an ATAC library. The product is taken through a pre-amplification PCR step to fill gaps and ensure maximum recovery of barcoded ATAC fragments. Subsequently, the pre-amplified product is used as input for ATAC library construction. Resulting libraries were sequenced using 150bp paired-end reads on the Illumina NovaSeq 6000 system at a recommended depth of 25,000 read pairs per cell. Following sequencing, BCL or

FASTQ files were analyzed using the Cell Ranger ATAC analysis pipeline. Cell Ranger ATAC performs sample demultiplexing, barcode processing and identification of open chromatin regions in single cells. Secondary analyses, such as dimensionality reduction, cell clustering, and peak analysis were performed through Seurat and detailed bellow (Signac Pipeline[2]).

**CFU assay.** To assess clonogenic progenitor frequencies, $3 \times 10^4$ CD34+ HSPC cells were plated in methylcellulose containing SCF, GM-CSF, IL-3 and EPO (H4434; STEMCELL Technologies). Colonies were scored 14 days later. Experiment was performed in triplicates.

**Zero-inflated CpG filtering for methylation analysis**

To filter Zero inflated CpGs inherent in such sequencing-based protocol, we filter CpG loci according to their detection rate across samples and the Msp1 reference. To optimize the threshold of filtering, we created two global quality score of the methylation data: one based on the percentage of real non-zeros hypermethylation across highly methylated loci, and the other representing the dependency of the sample covariance to the percentage of detected loci by samples. The threshold for each CpG quality metric were chosen according to the knee of the distribution curve that represents the increase of the global quality scores across the tested thresholds. Accordingly, we filtered out CpG loci with less than 5 Msp1 count (over 76,541,158 total read counts), with a Confidence Score (defined as the sum of all count for HpaII libraries and Msp1 library, normalized by library size) less than 16/100M reads, and with more than 95% of samples with zero count. We also removed CpG loci when presenting at least 70% of zero count across samples and without any sample having a methylation score between 0 and 10 (excluded). 754,931 out of 1,709,224 CpGs were conserved for further analysis.

**Linear modeling and differential methylation analysis**

To identify differentially methylated CpGs (DMCs), we performed linear regression and statistical modeling using the Limma R package.[3] To identify confounders to be included in the model, we assess the linear correlation between technical and biological covariates and the 10 first principal components (PCs) computed from the DNA methylation data (**Supplemental Figure 1**). PC1 was associated with Group (LGA or CTRL), Sex, Maternal age, Ethnicity (latino or not), and Cohort (from already published batch or from the new batch), so we included them in the model. However, the main factor of covariability was the detection rate (number of loci detected in each sample). This variable not only depends on technical variability, but also have a biological component (fully methylated CpGs would not be detected, so a global DNA hypermethylation will results on a low detection rate). Therefore, to preserve the biological influence while isolating the technical variability of the detection rate, we classified each sample within each group in four equally sized classes from “very low detection rate” to “very high detection rate”. Contrary to the detection rate, this detection level within group was not correlated with the biological covariates (*e.g.*, group and the maternal age) thus we included it in the linear model as well. We also included PC2 to the model as it was the second contributor to variability in our dataset and was not correlated to any known covariates. We performed a differentially methylated analysis (LGA vs CTRL) considering the new batch of samples counting 16 CTRL and 16 LGA as well as considering all samples together adding to the newly generated data the already published data, for a total number of samples of 34 CTRL and 36 LGA. 2 CTRL samples from previously generated data were excluded for this analysis as ethnicity information were missing and was part of our linear model.

**Gene-methylation score**

*LinkWeight.* We link CpG to gene following one of these 2 approaches: i) CpG were linked to gene based on transcription start site (TSS) distance, in respect to a 200kb window. If more than one gene is located at +/- 200 kb of the CpG locus, we kept only the closest gene based on its TSS.

For this TSS based link, the LinkWeight was defined as:

If $x < 1$ kb: LinkWeight = 1

If $x < 20$ kb: LinkWeight = $0.5 + 0.5 \times \sqrt{1000/x}$

Else: LinkWeight = $0.5 \times \sqrt{1000/x}$

Where x is the CpG distance to the TSS

or ii) CpG were linked to gene based on expression quantitative trait loci (eQTL) data, if located in a 1 kb window around associated single nucleotide polymorphism (SNP). We used blood specific and tissue wide SNPs-gene cis-association from the Genotype-Tissue Expression (GTEx) analysis V8 (dbGap Study Accession: phs000424.v8.p). 1.2M out of a total of 1.7M of CpGs were associated to genes including 320K associated thanks to GTEx data.

For this SNP based link, the LinkWeight was defined as:

If $\mathrm{mean}(-\log10(p_{snp})) > q90$: LinkWeight = 1

Else: LinkWeight = $\mathrm{mean}(-\log10(p_{snp}))/q90$

Where $p_{snp}$ is the nominal *p*-value of the association between the SNP and the gene; q90 is the 90th percentile of the *p*-value distribution.

*RegWeight.* To give more weight to CpG methylation change that have more chance to impact gene expression, we weighted each CpG methylation change according to the CpG location in candidate or known regulatory regions. This RegWeight is defined as 0.5 + 1.5 × (ChIPScore + EnsRegScore) / 2 where ChIPScore refers to CD34+ specific genomic annotation defined using CD34+ specific histone marks as previously described[4] and EnsRegScore refers to regulatory regions defined based on the Ensembl Regulatory build hg19 genome annotation.[5]

The ChIPscore is defined as:

If l in Enhancer or promoter region: ChIPScore = 1

If l in poised Enhancer region: ChIPScore = 0.75

If l in Gene-body region: ChIPScore = 0.5

If l in HeteroChromatin region: ChIPScore = 0

Where l is the CpG locus.

The EnsRegScore is defined as:

If l in CTCF Binding Site, Promoter, Enhancer: EnsRegScore = 0,5

If l in Open chromatin, Promoter Flanking Region: EnsRegScore = 0,25

If l in TF motif binding site region: EnsRegScore = EnsRegScore + 0,5

Where l is the CpG locus.

To optimize the LinkWeight and RegWeight, a simulated dataset of 36,720 CpGs that recapitulates the range of the different methylation metrics (methylation change and *p*-value of the significance) and genomic features (location in regulatory region, distance to TSS, presence in eQTL region) present in our original dataset was created. This dataset was then used to scale weights to respect the following importance order: methylation change = pvalue > TSS distance = eQTL region > CD34+ specific genomic annotation > Ensembl Regulatory build annotation.

2) To concatenate CpG-Scores at gene level: gene-methylation score

To summarize the CpG methylation change at the gene level, we aggregated the CpG-Scores into a gene-methylation score by taking care of i) alleviate the arbitrary number of CpGs per gene and ii) interpret differently CpG influences located on the promoter of them in others genomic region.

To alleviate the influence of the number of CpGs linked to a gene, the $Weight_{nCpG}$ defined above was optimized by modelling the influence of each CpGs and gene features on the gene-methylation score and empirically test different $Weight_{nCpG}$ parameters to select the parameter that will preserve the influence of key factors (methylation change, significance of the methylation change, genomic context) but correct for the number of associated CpGs. This model was tested using the linear regression function from base R function and was defined as followed:

```
gene_score~n.cpg.gene+n.cpg.sig.gene+pval+meth.change+chromatin_feature+ensembl_reg_score+in_eQTL_region+abs(tss_dist))
```

Where for each gene, `n.cpg.gene` is the number of CpG linked and `n.cpg.sig.gene` is the number of CpGs significantly differentially methylated (*p*-value<0.001). `pval` and `meth.change` are the p-value and $log_2$(FoldChange) of the methylation change, `chromatin_feature` is the CD34+ specific genomic annotation, `ensembl_reg_score` is the Ensembl Regulatory build annotation, `in_eQTL_region` is the presence of the CpG in eQTL region, and `abs(tss_dist)` is the absolute distance between the CpG and the TSS.

#### Gene-methylation score validation

To validate that the gene-methylation score better highlights genes susceptible to be transcriptionally impacted by the methylation change than others methylation metrics, the association between these methylation metrics and gene expression change was tested. To do that, DEGs (adjusted p-value <0.05) found in 6 LGA vs 8 Control HSPCs samples using the pseudo-bulk DESeq2 analysis was used as response variable. Then, Wilcoxon tests was performed for each methylation metrics on the difference of this methylation metrics between DEGs and non-DEGs. The methylation metrics tested were: *p*-value, $-\log_{10}(p\text{-value})$, $\log_2(\text{FoldChange})$ and $\log_2(\text{FoldChange})*-\log_{10}(p\text{-value})$ of the most significant CpG for each gene, as well as, average of the *p*-value, the $-\log_{10}(p\text{-value})$, the $\log_2(\text{FoldChange})$, and the $\log_2(\text{FoldChange})*-\log_{10}(p\text{-value})$ across all CpGs link to the gene. Considering the 9 methylation metrics tested, the gene-methylation score presents the best association with DEGs (**Figure 2**).

#### Gene Set enrichment analysis

We assessed enrichment for biological pathways in epigenetically altered genes by performing gene set enrichment analysis (GSEA). We ranked the genes based on their methylation gene-score, from the most epigenetically altered (highest gene-methylation score) to the less epigenetically altered (lowest gene-methylation score) and performed GSEA using the clusterProfiler package.[6] For KEGG pathways and Gene Ontology (GO) terms, we used the gseKEGG and gseGO functions, respectively. For GWAS physiological or pathological traits, we obtained the list of susceptibility genes for each trait from the “reported genes” column of the GWAS catalog database[7] and used the GSEA function. We excluded for traits with less than ten susceptibility genes. For Regulon enrichment in epigenetically altered genes or differentially expressed genes (see Coregulatory network analysis section), we also used this GSEA function using Regulon as gene-set considering the gene-methylation score and the $log_2FC$ x ($-log_{10}$($p$-value)) for the expression.

**Transcription factor motif enrichment analysis**

We performed transcription factor (TF) motif enrichment analysis using the HOMER software.[8] We assess the enrichment for 437 known TF motifs (included in the HOMER software) in ±20 bp regions around the DMCs ($p$-value < 0.001 and |DNA methylation change| > 25) compared to 50K randomly selected loci (background) from our dataset presenting a high GC content.

**Single cell RNA-seq dataset preprocessing**

Unique Molecular Index (UMI) Count Matrices for gene expression and for HTO libraries were generated using the CellRanger count (Feature Barcode) pipeline. Reads were aligned on the GRCh38-3.0.0 transcriptome reference (10x Genomics). Filtering for low quality cells according to the number of RNA, genes detected, and percentage of mitochondrial RNA was performed. For HTO sample, we normalized the HTO matrix using centered log-ratio (CLR) transformation and cells was assigned back to their sample of origin using HTODemux function of the Seurat R Package (v4).[9] Then, we normalized the gene expression matrix for cellular sequencing depth and regress for mitochondrial percentage and cell cycle phases differences using the variance stabilizing transformation (vst) based Seurat::SCTransform function. **Supplemental Table 8** contains information on number of cells per sample.

**Early Hematopoietic reference map (Hematomap) creation and mapping**

We integrated the 7 CTRL datasets ($n$ = 16,912 cells) with the Seurat (v4) Canonical Correlation Analysis (CCA) and graph based integration tool using the 3,000 most expressed genes across datasets to correct for batch effect. The 30 first dimensions of the PCA of the batch effect corrected matrix was used to generate the Shared Nearest-neighbor (SNN) graph and the UMAP. Graph-based clustering using Louvain algorithm with a resolution parameter of 0.6 on the FindCluster function was used to cluster cells. Each cluster was annotated using cell type specific markers. Markers for each cluster were identified using FindAllMarkers function with default parameter. Genes were then ranked based on their expression fold change the difference of detection of this gene in the cluster versus all other clusters and the specificity for the cluster, and top cluster-specific genes were compared with published cell type-specific genes. This hematomap was then used as reference to annotate all datasets for the different hematopoietic cell types thanks to Seurat (v4) MapQuery function.

**PseudoBulk differential expression analysis**

PseudoBulk differential expression analysis between LGA and CTRL cells within each hematopoietic cell type was performed using the DESeq2 R package[10] to assess influence of group, stimulation and interaction between group and stimulation. We aggregated the gene count by sample within each cell type before performing the differential expression analysis. We included the batch and the sex as cofounding covariates in the negative binomial Generalized linear model (GLM).

**Over-representation test**

Over representation test was performed on Differentially Expressed Genes (DEGs) using the enrichGO and enrichKEGG of the clusterProfiler R package depending on the Gene sets of interest.

**Population distribution analysis**

To test for cell type proportion difference between CTRL (*n* = 6) and LGA (*n* = 6) samples we used the Wilcoxon rank-sum test for comparing the proportions of each cell type between in LGA and Control samples. Two samples were assigned as outliers based on lineage distribution using the boxplot, *i.e.*, Tukey method, and therefore excluded from this analysis.

**Pseudotime analysis**

Differentiation trajectory analyses were conducted with monocle[11] (https://www.bioconductor.org/packages/monocle/). Preprocessed Seurat object were imported unsing *importCDS* function from the monocle R package. Monocle's *orderCells* function was used to arranged cells along a pseudo-time axis to indicate their position in a developmental continuum. Monocle generates for each cell a pseudotime value in respect to predefined cell of origins (roots). Here the same LT-HSC cells were used as roots for the whole integrated dataset to have comparable pseudotime across conditions. We specify the root of the trajectory programmatically, as recommended, by first grouping the cells according to which trajectory graph node they are nearest to. Then, calculating what fraction of the cells at each node come from the earliest time point. Then by picking the node that is most heavily occupied by early cells and returns that as the root.

To test the difference in pseudotime between LGA and CTRL samples, we modelled the influence of LGA on the pseudotime at the cell level with a Linear Mixed-Effects Model using lme4 R package and estimated the p-value using the lmerTest R package. The model formula used was:

pseudotime ~ 1 +group+ group:lineage_hmap + (1|sample)

Where pseudotime is the pseudotime of the cell, group is the group (LGA or Control), group:lineage_hmap is the interaction between the group and the lineage of the cell, and sample is the sample from which the cell comes from. To test if the LGA conditions was associated with a reduced proportion of cells with low pseudotime (undifferentiated state), Wilcoxon test was performed on the percentage of cells below each pseudotime from 0 to 30. The outliers identified in the population distribution analysis was also excluded for this analysis

**Coregulatory network analysis**

To identify coregulated genes by a same TF (regulons) on our dataset, we used the SCENIC workflow[12] on a batch corrected matrix for all samples (CTRL, CTRL HTO and LGA HTO). The batch corrected matrix was obtained using the Seurat (v4) integration tool. We used the GENIE3 R package to identify co-expressed gene modules and the RcisTarget R package to infer potential TF targets for each module. Regulatory modules (regulons) were identified from co-expression and DNA motif analyses. Regulons were then evaluated in each cell to ascertain their activities by the AUCell package in Bioconductor. To reduce the computational time during the coexpression module identification using the GENIE3 algorithm, we subset the batch corrected gene expression matrix by picking randomly 100 cells by sample and by cell type. After regulons identification based on this subset of cells, we score the regulons activity in each cell on the entire dataset using the AUCell algorithm. All TFs and targeted genes of high confidence regulons based on RcisTarget was used to build a directed graph representing interaction between TF and genes.

**Single cell ATAC-seq data processing**

Reads were aligned to the GRCh38 reference and peaks were called using Cell Ranger ATAC pipeline (10X Genomics) generating a unified peak count matrix. Cells with less than 500 counts in these peaks were filtered out. Overlapping peaks from the different libraries were merged to generate a unified set of peaks. Unified peaks with a width over 10kb or less than 20bp were filtered out.

Data were then analyzed through the Signac workflow[2]. EnsDb.Hsapiens.v86 annotation package with UCSC hg38 style was used to annotate peaks. For QC filtering, cells with less than 5000 or above 60000 counts in peaks as well as cells with more than 15% of reads outside peaks regions were filtered out. Cells with more than 0.15% of reads in blacklist region as defined by the ENCODE project[13] were also filtered out. Finally, cells with a ratio of mononucleosomal to nucleosome-free fragments (nucleosome signal) over 1 or a TSS enrichment less than 2 or more than 10 were filtered out. Before clustering, the peak count matrix was normalized using term frequency-inverse document frequency (TF-IDF) to correct for differences in cellular sequencing depth and give higher values to more rare peaks. Singular value decomposition (SVD) was used to reduce dimensions based on latent semantic indexing (LSI) approach. Batch effect was corrected from these LSI components using RunHarmony function[14] default parameters. First LSI component was excluded from downstream clustering steps, as this component was highly correlated to the sequencing depth (technical variability). UMAP and graph-based clustering using Share Nearest Neighbour Graph (SNN) and smart local moving (SLM) clustering algorithm was performed on the 2nd to 30th batch corrected LSI components. Cells were then annotated for hematopoietic cell type using a label transfer approach based on the hematomap reference. Gene level count matrix was first generated using GeneActivity function and normalized using SCTransform. Cells anchors between hematomap and ATAC datasets were defined thanks to FindTransferAnchors function based on canonical correlation analysis (CCA) reduction. Hematopoietic lineage labels were then predicted for each ATAC cells using these anchors and the TransferData function.

To identify putative lineage specific peaks, peak calling was then performed at lineage level using MACS2 based CallPeaks function and the predicted lineage label as grouping variable. Lineage specific peaks were identified using FindMarkers function (Signac) with Logistic Regression (LR) models including cellular sequencing depth as latent variable.

**Supplemental Table 8** contains information on number of cells per sample.

**Differential accessibility analysis and TF motif enrichment analysis**

Peaks differentially accessible between two specific conditions were identified using FindMarkers function (Signac) with Logistic Regression (LR) models including cellular sequencing depth as latent variable.

FindMotifs function (Signac) was used to calculate enrichment for TF motif in specific set of peaks compared to a background set of peaks. For TF motif enrichment in lineage specific peaks, the default parameters were used (background correspond to 40000 peaks representative of sequence characteristics of the query features). For TF motif enrichment in DMCs containing peaks, background was defined as all peaks containing methyl assay queried CpGs.

#### Integrative Gene Regulatory Network construction

To build the gene regulatory network integrating gene expression and open chromatin data, we used TF-target interactions from the SCENIC regulons analysis. TF-target interactions were included if TF motif was present in peak and peak was associated to HSC. We represented TF-target interactions four our TFs of interest: EGR1, KL2 and KLF4 using the *network* R package. Node represents gene of the network and was annotated as following: i) if the gene is a TF and if the gene is differentially expressed in HSC comparing LGA vs CTRL (adjusted *p*-value < 0.05 and |fold change| >0.5). Gene label represent Gene-methylation score. Edge represent TF-gene target regulatory link and was annotated as following: if the peak linking the TF to the target gene (by presence of TF motif in it) i) have DMCs comparing LGA vs CTRL (p-value <0.001 and |methylation difference| > 25), ii) are differentially accessible comparing LGA vs CTRL HSC (adjusted p-value<0.001 and $|\log_2 FC|>0.25$).

Supplemental Figure 1

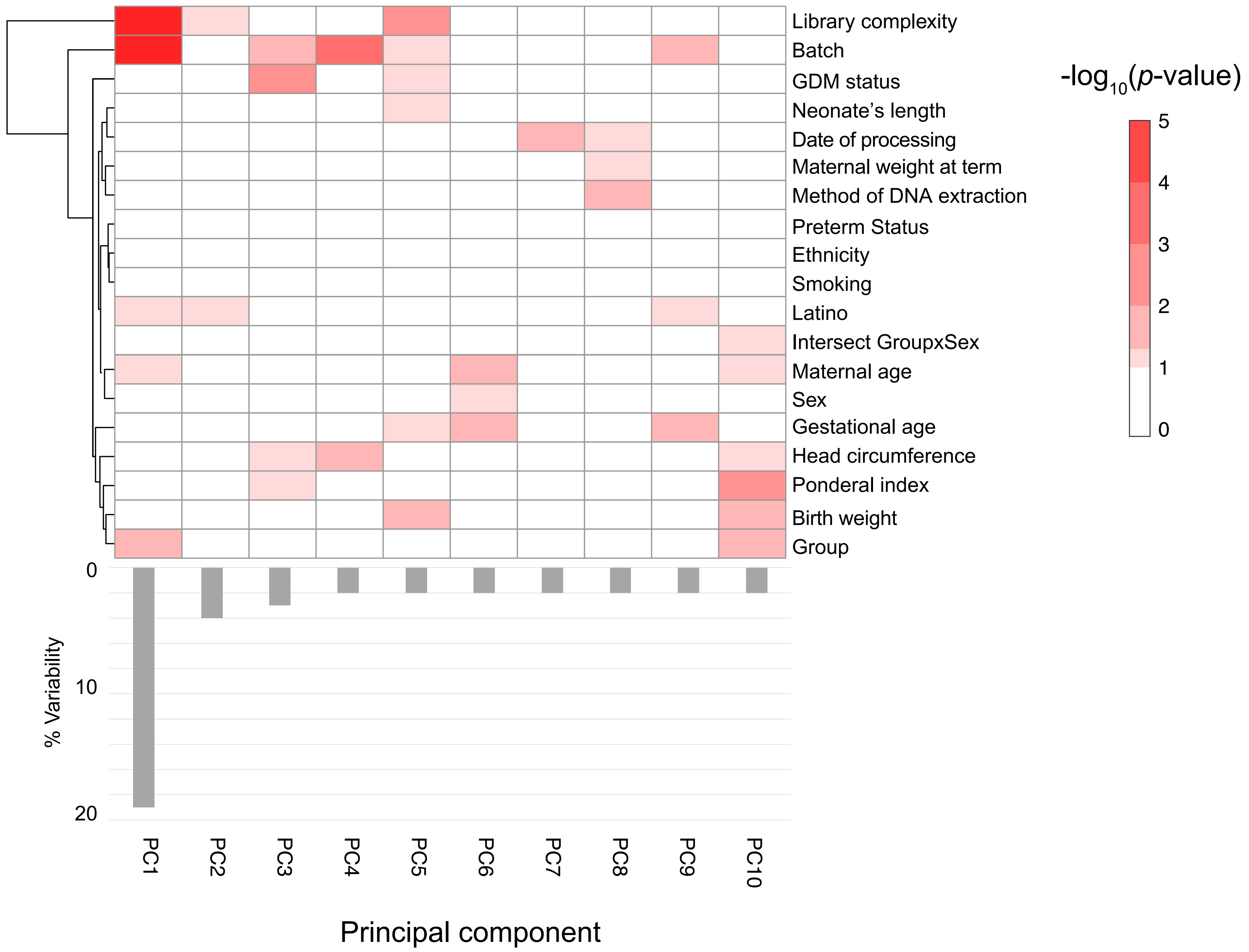

Supplemental Figure 2

A

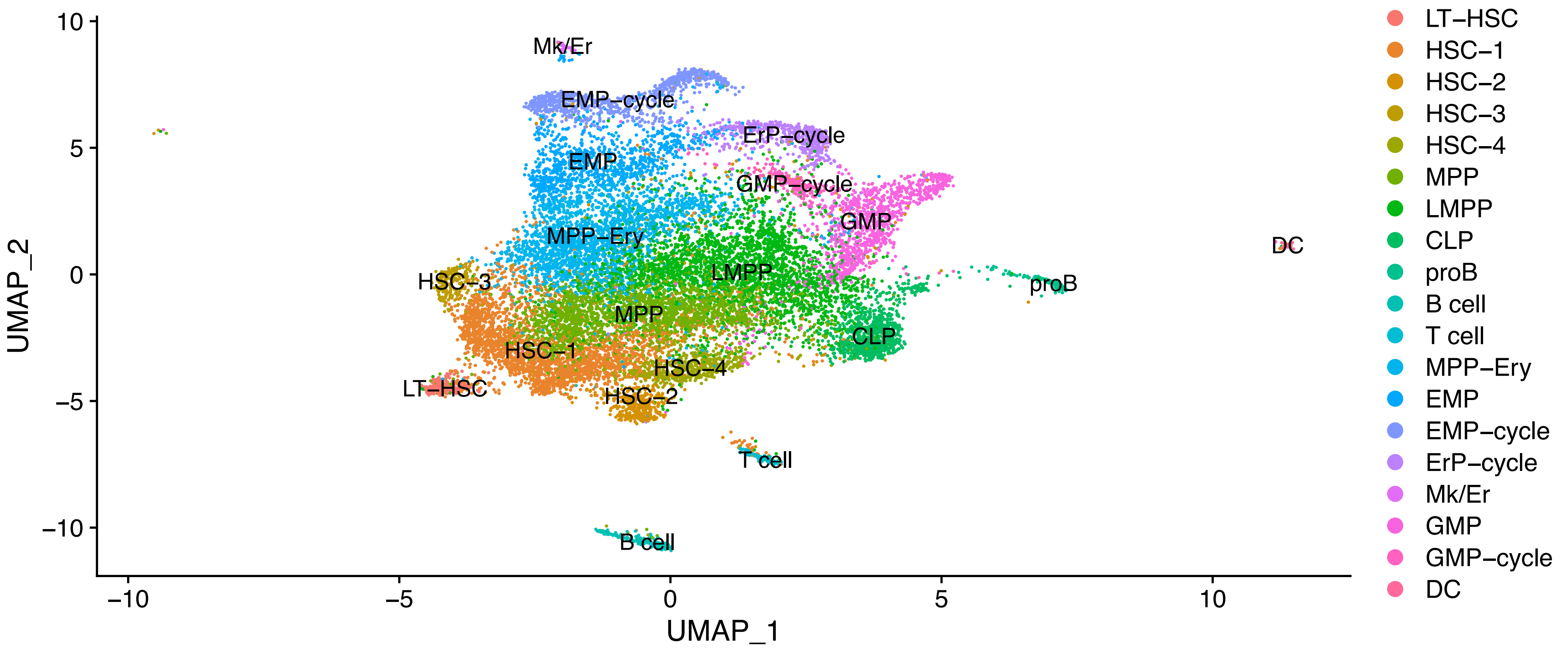


B

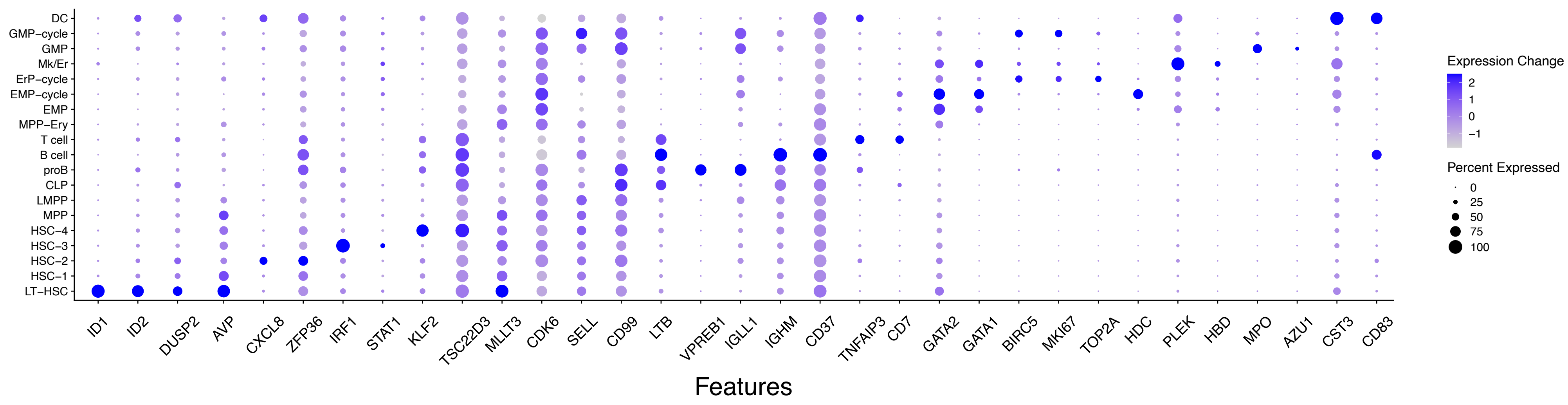

Supplemental Figure 3

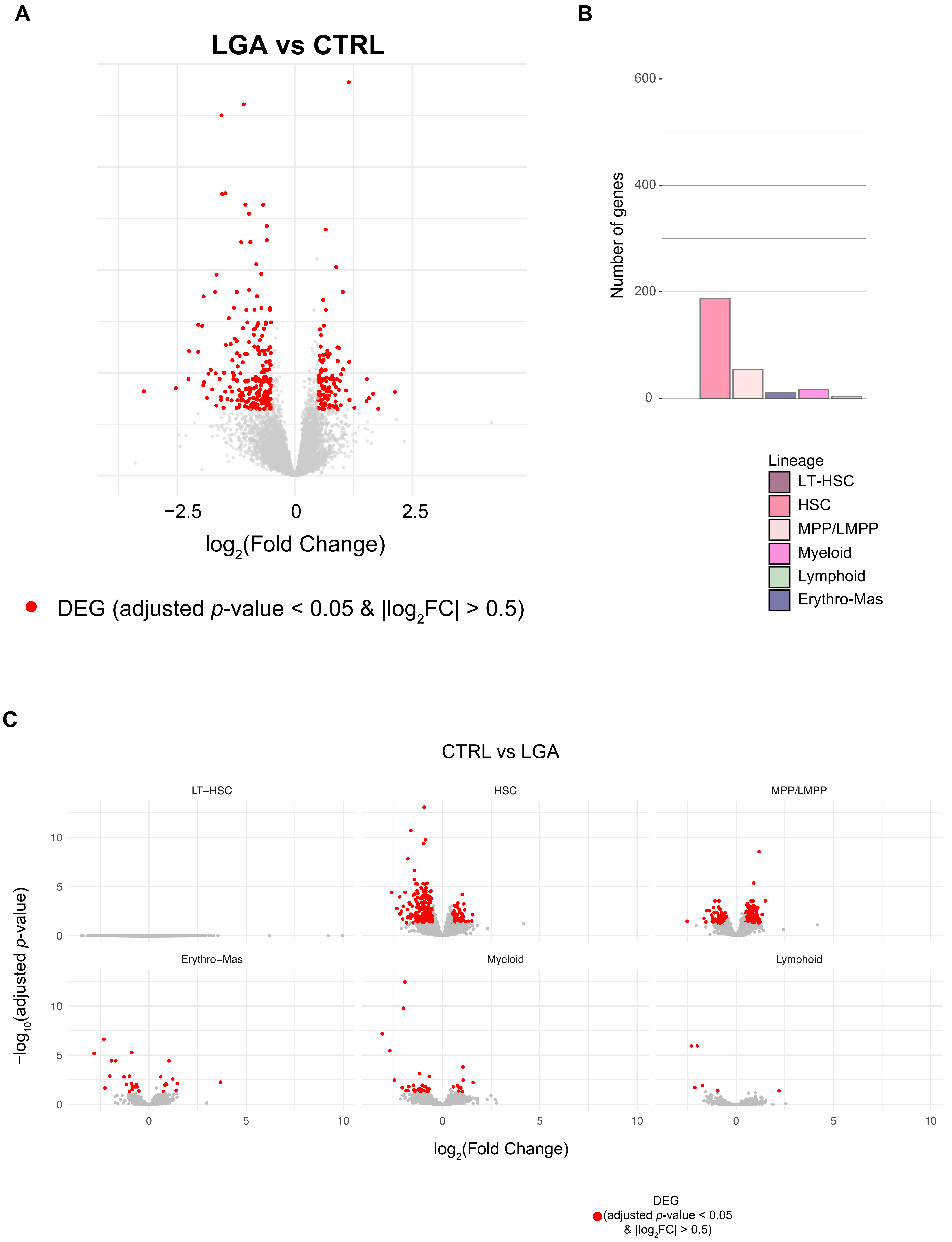

### Supplemental Figure 4

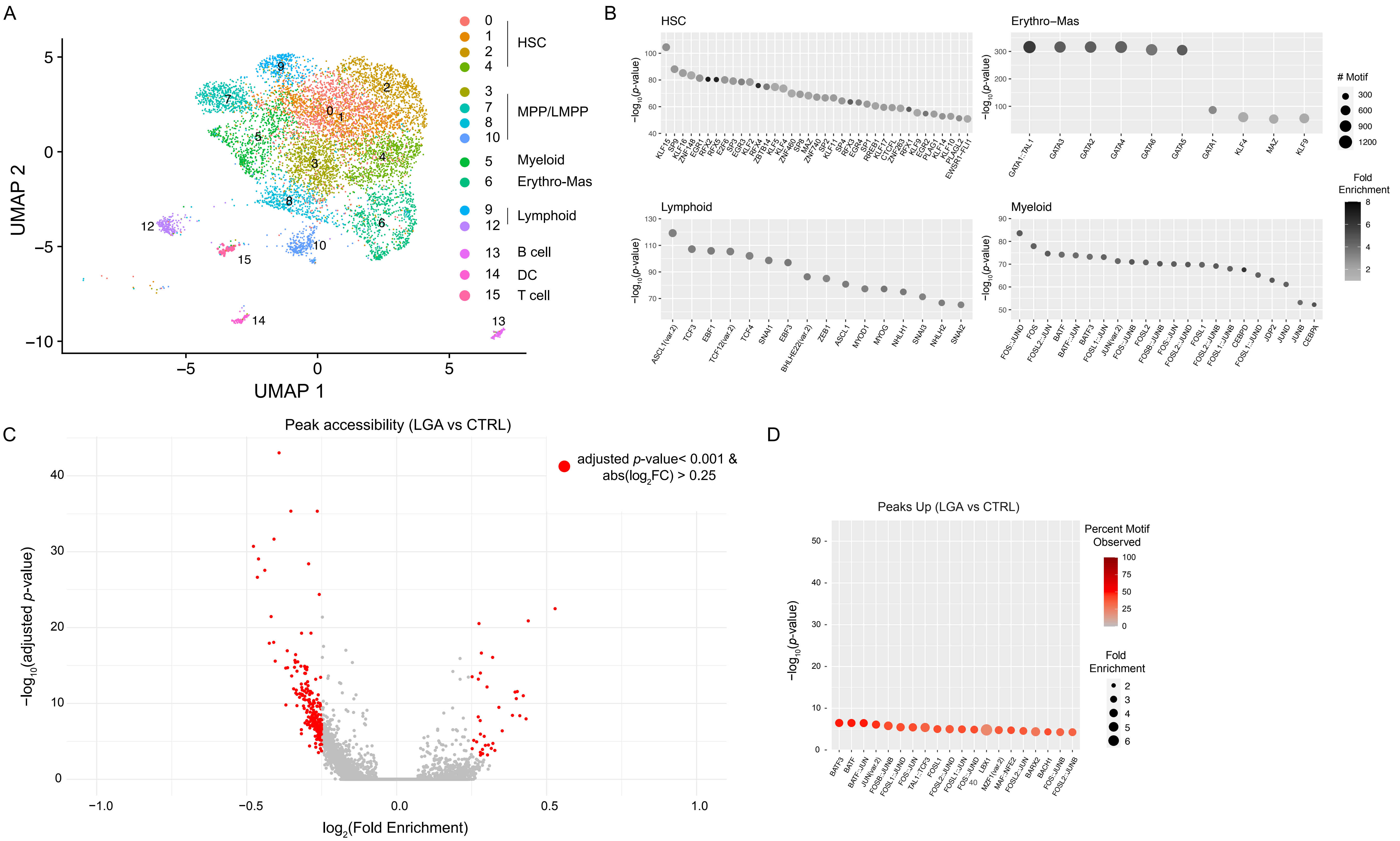

Supplemental Figure 5

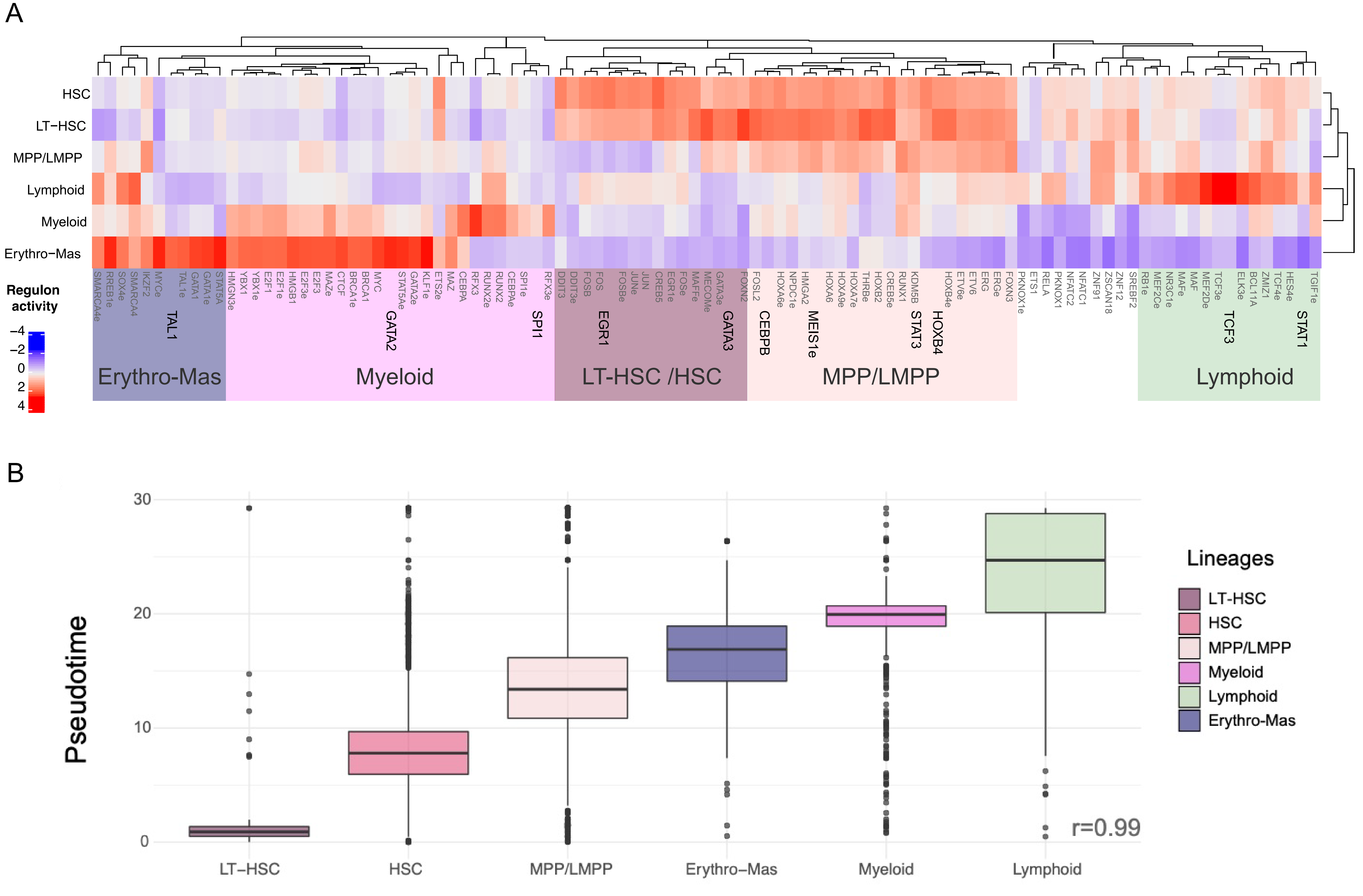
